# Knomics-Biota - a system for exploratory analysis of human gut microbiota data

**DOI:** 10.1101/274993

**Authors:** Anna Popenko, Alexander Tyakht, Anatoly Vasilyev, Ilya Altukhov, Daria Efimova, Vera Odintsova, Natalya Klimenko, Robert Loshkarev, Maria Pashkova, Anna Elizarova, Viktoriya Voroshilova, Sergei Slavskii, Yury Pekov, Ekaterina Filipova, Tatyana Shashkova, Evgenii Levin, Dmitry Alexeev

## Abstract

**Summary:** Metagenomic surveys of human microbiota are becoming increasingly widespread in academic research as well as in food and pharmaceutical industries and clinical context. Intuitive tools for exploration of experimental data are of high interest to researchers. Knomics-Biota is a Web-based resource for exploratory analysis of human gut metagenomes. Users can generate analytical reports that correspond to common experimental schemes (like case-control study or paired comparison). Statistical analysis and visualizations of microbiota composition are provided in association with the external factors and in the context of thousands of publicly available datasets.

**Availability and Implementation:** The Web-service is available at https://biota.knomics.ru.

*Contact:*.

*Supplementary information:* Supplementary figures are available at *Bioinformatics* online.

## Introduction

The last decade was marked by an explosive growth of experimental data characterizing human-associated microbial communities using metagenomic approach. Previously employed mainly by the academic community, now metagenomics are used in the industry to assess structure, functions and dynamics of microbiota composition - particularly, to identify the impact of products and medical drugs on human microbiota and health. Of particular importance are visual and statistical exploration of important functions of microbiota (like antibiotic resistance (Yarygin et al., 2017) and dietary fiber catabolism (Yarygin et al., 2017)) in the global context of publicly collected data. Lower costs and increasing popularity make metagenomics further available to smaller companies and research facilities that often lack a dedicated staff bioinformaticians that can perform expert statistical analysis and visualization according to state-of-art guidelines (Odintsova 2017), (Sudarikov 2017). In order to optimize the conversion of metagenomic surveys results into biomedically important knowledge and advance the global progress in microbiota research, we developed Knomics-Biota, a Web-service for metagenomic data analysis that allows users without advanced skills in bioinformatics and software development to turn their “raw” data” into intuitive analytical reports. The datasets can be accompanied with metadata that can include, besides external factors like age and clinical status, the factors related to experimental design - distribution between case and control groups, paired correspondence of the samples, etc. After automatic analysis in the cloud, user is provided with analytical reports describing the results of metagenomic analysis - from data quality check and composition profiles to statistical hypothesis testing. Interactive visualization modules allow to explore the interactions between microbiota and factors in details and yield novel biological hypotheses. Analysis of metabolic potential includes manually curated pathways reflecting gut microbiota functions highly relevant for human health - like synthesis of short-chain fatty acids (SCFAs) and vitamins.

## Knomics-Biota functions and use examples

The computational backend of the system is located in the cloud (see Supplementary Figure 1) and makes use of publicly available software solutions. The front-end interface of the web service is implemented using Yii framework, and interactive visualisations make use of d3js library. After signing up, a user can create a project in his/her account and upload the “raw” data - metagenomic reads in FASTQ format obtained via amplicon (16S rRNA) or “shotgun” sequencing.

**Figure 1.**
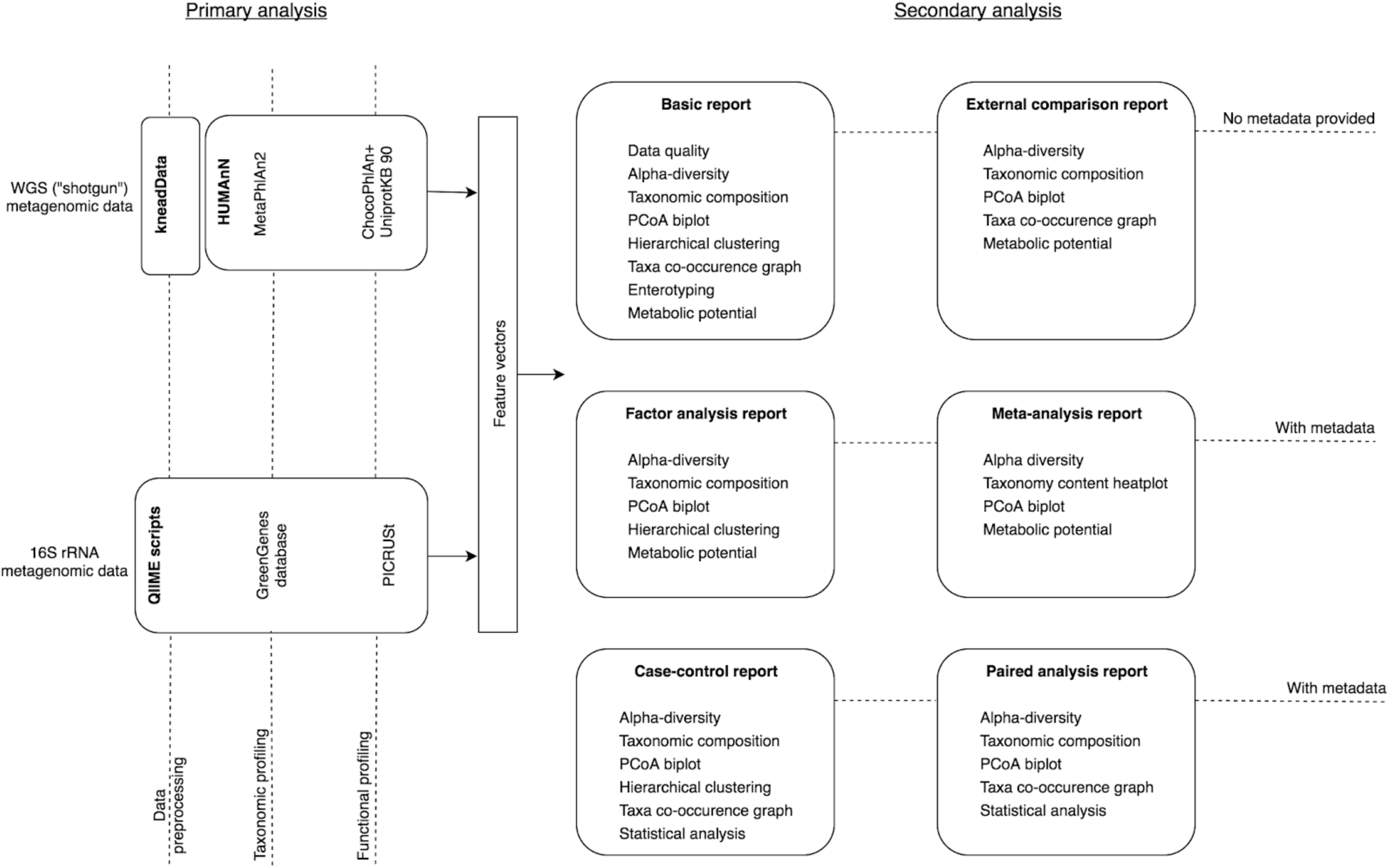
Workflow of the Knomics-Biota web-service.

General logic of Knomics-Biota service includes two components: primary and secondary analysis (see Fig. 1). Primary analysis component encompasses basic processing of the reads to obtain microbiota composition profiles. For each of the 16S rRNA and “shotgun” formats, primary analysis component produces feature vectors including relative abundance of microbial taxa at various ranks as well as of gene groups and metabolic pathways according to KEGG Orthology and Enzyme Commission (EC) nomenclatures. Additionally, some functions are analyzed in a dedicated way due to their importance for human health - synthesis of vitamins and SCFAs. These functions are assessed for each sample using curated pathways (see Supplementary Figure 2).

Secondary analysis component implemented using Python v. 3.2 performs statistical analysis of these feature vectors (together with the metadata, if provided) and generates static figures as well as input for interactive visualization modules. The workflow of the secondary analysis varies depending on the choice of report type study by the user (see Fig. 1).

The basic report is performed essentially for any user data. It includes quality check of “raw” data, assessment of relative abundance of taxa and gene groups as well as alpha-diversity. Hierarchical clustering, enterotyping (Arumugam et al., 2011) and metabolic potential prediction are performed. Besides basic visualizations, interactive modules are provided including heatmap, PCoA plot, alpha-diversity plot and co-occurrence network (Kurtz et al., 2015).

The case-control report is available when the corresponding metadata is provided. The functionality of the interactive modules is extended to allow comparison of the microbiota composition between the two groups. Statistical analysis is performed to identify the respective significant differences. Besides basic composition features, gut microbiota-specific characteristics of interest are evaluated and compared between the groups: these include metabolic potential for synthesis of vitamins and SCFAs.

Paired analysis report has a workflow that is similar to a case-control scenario but is performed to compare paired samples (for instance, those collected from the same subjects before and after the antibiotic consumption). The details of interactive modules and statistical analysis are adapted, respectively.

In the external comparison report, the whole set of user metagenomes can be compared with any of the publicly available metagenomic datasets from tens of scientific publications that have been precomputed, annotated and stored in Knomics-Biota database. User can select a subset of these data by subjects’ disease or other factors.

Factor analysis report is run if metadata provided: the service performs multifactor analysis to identify significant associations between microbiota composition and external factors (age, BMI, clinical status, etc.). The interactive modules are adapted to include controls over the display of these factors.

Meta-analysis report is a combination of external comparison report and factor analysis reports: it allows to assess the links between microbiota and factors over the union of user and external datasets.

When a user wants to analyze series of consecutive metagenomic profiles (e.g. multiple points collected from the same individual during the course of dietary intervention), one can use a time series report to explore the data using adapted visualizations and perform specific statistical analysis like temporal stability evaluation.

## Conclusions

Knomics-Biota service is a convenient tool for exploratory analysis of metagenomes in the context of publicly available data. Besides gut microbiota, the system is ready for processing metagenomes from an arbitrary environment allowing users with and without expertise in bioinformatics to gain insights into system biology of complex microbial communities.

